# Expanded Phylogenetic Diversity and Metabolic Flexibility of Microbial Mercury Methylation

**DOI:** 10.1101/2020.01.16.909358

**Authors:** Elizabeth A. McDaniel, Benjamin Peterson, Sarah L.R. Stevens, Patricia Q. Tran, Karthik Anantharaman, Katherine D. McMahon

## Abstract

Methylmercury is a potent, bioaccumulating neurotoxin that is produced by specific microorganisms by methylation of inorganic mercury released from anthropogenic sources. The *hgcAB* genes were recently discovered to be required for microbial methylmercury production in diverse anaerobic bacteria and archaea. However, the full phylogenetic and metabolic diversity of mercury methylating microorganisms has not been fully explored due to the limited number of cultured, experimentally verified methylators and the limitations of primer-based molecular methods. Here, we describe the phylogenetic diversity and metabolic flexibility of putative mercury methylating microorganisms identified by *hgcA* sequence identity from publicly available isolate genomes and metagenome-assembled genomes (MAGs), as well as novel freshwater MAGs. We demonstrate that putative mercury methylators are much more phylogenetically diverse than previously known, and the distribution of *hgcA* is most likely due to several independent horizontal gene transfer events. Identified methylating microorganisms possess diverse metabolic capabilities spanning carbon fixation, sulfate reduction, nitrogen fixation, and metal resistance pathways. Using a metatranscriptomic survey of a thawing permafrost gradient from which we identified 111 putative mercury methylators, we demonstrate that specific methylating populations may contribute to *hgcA* expression at different depths. Overall, we provide a framework for illuminating the microbial basis of mercury methylation using genome-resolved metagenomics and metatranscriptomics to identify methylators based upon *hgcA* presence and describe their putative functions in the environment.

**IMPORTANCE:** Specific anaerobic microorganisms among the *Deltaproteobacteria, Firmicutes*, and *Euryarchaeota* have been shown to produce the bioaccumulating neurotoxin methylmercury. Accurately assessing the sources of microbial methylmercury production in the context of phylogenetic identification, metabolic guilds, and activity in the environment is crucial for understanding the constraints and effects of mercury impacted sites. Advances in next-generation sequencing technologies have enabled large-scale, cultivation-independent surveys of diverse and poorly characterized microorganisms of numerous ecosystems. We used genome-resolved metagenomics and metatranscriptomics to highlight the vast phylogenetic and metabolic diversity of putative mercury methylators, and their depth-discrete activities in the environment. This work underscores the importance of using genome-resolved metagenomics to survey specific putative methylating populations of a given mercury-impacted ecosystem.

## INTRODUCTION

Methylmercury is a potent neurotoxin that biomagnifies upward through food webs. Inorganic mercury deposited in sediments, freshwater, and the global ocean from both natural and anthropogenic sources is converted to the bioavailable and biomagnifying organic methylmercury (1, 2). Methylmercury production in the water column and sediments of freshwater lakes is one of the leading environmental sources that leads to fish consumption advisories (3, 4). Additionally, the majority of methylmercury exposure stems from contaminated marine seafood consumption (5). Consumption of methylmercury-contaminated fish is the most problematic for childbearing women and results in severe neuropsychological deficits when developed fetuses are exposed (6, 7). The biotransformation from inorganic to organic mercury is thought to be carried out by specific anaerobic microorganisms, namely sulfur-reducing (SRB) and iron-reducing (FeRB) bacteria and methanogenic archaea (8–10). Recently, the *hgcAB* genes were discovered to be required for mercury methylation. The *hgcA* gene encodes a corrinoid-dependent enzyme that is predicted to act as a methyltransferase, and *hgcB* encodes a 2[4Fe-4S] ferredoxin that reduces the corrinoid cofactor (11). Importantly, these genes are known to only occur in microorganisms capable of methylating mercury. This discovery has allowed for high-throughput identification of microorganisms with the potential to methylate mercury in diverse environmental datasets based upon *hgcAB* sequence presence (12, 13).

Currently, the known diversity of mercury methylating microorganisms has been described based on cultured isolates, amplicon sequencing of the 16S rRNA and/or *hgcA* loci of mercury impacted sites, *hgcA* quantitative PCR (qPCR) primers targeting specific groups, or identification of *hgcA* on assembled metagenomic contigs (11, 13–20). However, the complete phylogenetic and metabolic diversity of microorganisms capable of methylating mercury is not fully understood due to previous efforts focusing on a few cultured representatives, and poor predictive power of methylation potential based on 16S rRNA phylogenetic signal (8, 10, 21). Additionally, amplicon sequencing is limited in detection of phylogenetic diversity, and the metabolic capabilities of these microorganisms cannot be reliably studied using these techniques. Furthermore, a recent study of microbial methylation in sulfate-impacted lakes demonstrated that 16S rRNA amplicon-based surveys and *hgcA* clone libraries may not capture the true phylogenetic diversity of many methylating microorganisms (22).

In this study, we identified nearly 1,000 genomes containing the *hgcA* marker from publicly available isolate genomes, metagenome assembled genomes (MAGs), and novel bins assembled from three freshwater lakes. Using this collection of MAGs and reference genomes, we identified putative methylators spanning 30 phyla from diverse ecosystems, some of which from phyla that, to our knowledge, have never been characterized as methylating groups. The *hgcAB* phylogenetic signal suggests this diverse region originated through extensive horizontal gene transfer events, and provides insights into the difficulty of predicting methylation status using ribosomal sequences or “universal” *hgcA* primers (17, 22). Putative methylators span metabolic guilds beyond traditional sulfate and iron reducing bacteria and methanogenic archaea, carrying genes for traits such as such as nitrogen fixation transformations and metal resistance pathways. To understand *hgcA* expression in a potentially mercury impacted environment, we analyzed depth-discrete metatranscriptomes from a permafrost thawing gradient from which we identified 111 putative methylators. This work demonstrates the significance of reconstructing population genomes from mercury impacted environments to accurately capture the phylogenetic distribution and metabolic capabilities of key methylators in a given system.

## MATERIALS AND METHODS

### Accessed Datasets, Sampled Sites, and Metagenomic Assembly

We used a combination of publicly available MAGs, sequenced isolates, and newly assembled MAGs from three freshwater lakes. We sampled Lake Tanganyika in the East African Rift Valley, Lake Mendota in Madison, WI, and Trout Bog Lakes near Minocqua, WI. We collected depth-discrete samples along the northern basin of Lake Tanganyika at two stations (Kigoma and Mahale) (23). 24 samples were passed through a 0.2µm pore size-fraction filter, and used for shotgun metagenomic sequencing on the Illumina HiSeq 2500 platform at the Department of Energy Joint Genome Institute using previously reported methods with minor modifications (24). Each of the 24 metagenomes were individually assembled with MetaSPAdes and binned into population genomes using a combination of MetaBat, MetaBat2, and MaxBin. Across these binning approaches, approximately 4000 individual bins were assembled. To dereplicate identical bins assembled through different platforms, we applied DasTool (25) to keep the most representative bin for a given cluster, resulting in 803 total MAGs. Using a completeness/contamination cutoff of ≥70% and ≤10%, respectively, this resulted in approximately 431 MAGs. All dereplicated Lake Tanganyika genomes including identified putative methylators are available through the JGI IMG portal under GOLD study ID Gs0129147.

Five depth-discrete samples were collected from the Lake Mendota hypolimnion using a peristaltic pump through acid-washed Teflon tubing onto a 0.2µm Whatman filter, and used for Illumina shotgun sequencing on the HiSeq4000 at the QB3 sequencing center in Berkeley, CA, USA. Samples were individually assembled using MetaSPAdes (26), and population genomes were constructed using a combination of the MetaBat2, Maxbin, and CONCOCT binning algorithms. Resulting MAGs were aggregated with DasTool and dereplicated into representative sets by pairwise average nucleotide identity comparisons (25). Population genomes from samples collected from Trout Bog Lake were assembled as previously described (24, 27, 28). All Trout Bog Lake MAGs including the 4 putative methylators are available through the JGI IMG portal under GOLD study ID Gs0063444. All publicly available isolate genomes and MAGs were accessed from Genbank in August of 2019, resulting in over 200,000 genomes in which to search for *hgcA.* Genomes from Jones et al. 2019 were accessed from JGI/IMG in March of 2019 at GOLD study ID Gs0130353 (22). Only genomes of medium quality according to MiMAG standards with > 50% completeness and < 10% redundancy were kept for downstream analyses, as calculated with CheckM (29, 30).

### Identification of Putative Methylators

Using a collection of *hgcA* protein sequences from experimentally verified methylating organisms, we built a Hidden Markov Model (HMM) profile of the *hgcA* protein (11, 31). The constructed HMM profile was then used to identify putative mercury methylating bacteria and archaea. For all genome sequences, open reading frames and protein-coding genes were predicted using Prodigal (32). Using the HMM profile built for the *hgcA* marker, we searched all protein sequences with an e-value cutoff of 1e-50. Sequence hits with an e-value cutoff score below 1e-50 and/or a score of 300 were removed from the results to ensure high confidence in all hits. All archaeal and bacterial *hgcA* hits were concatenated and aligned with MAFFT (33). Alignments were manually visualized using AliView (34), and sequences without the conserved cap-helix domain reported in Parks et al. 2013 were removed (11).

From this set of putative methylators, the taxonomy of each genome was assigned and/or confirmed using both an automatic and manual classification approach. Each genome was automatically classified using the genome taxonomy database toolkit (GTDB-tk) with default parameters (35). Additionally, each genome was manually classified using a set of 16 ribosomal protein markers (36). If a genome’s taxonomy could not be resolved between the GTDB-tk classification and the manual ribosomal protein classification method, the genome was removed from the dataset (approximately 30 genomes). We kept specific genomes designated as “unclassified” as a handful of genomes with this designation belong to newly assigned phyla that have not been characterized in the literature, such as the proposed phyla BMS3A and Moduliflexota in the GTDB. This resulted in a total of 904 putative methylators used for downstream analyses as described. All genome information for each putative methylator including classification by each method, quality and genome statistics are provided in Supplementary Table 1 and summarized in Supplementary Figure 1.

### *hgcAB* and Ribosomal Phylogenies

We selected the highest quality methylator from each phyla (total of 30 individual phyla containing a putative methylator) to place in a prokaryotic tree of life. Bacterial and archaeal references (the majority from Anantharaman et al. (37)) were screened for the 16 ribosomal protein HMMs along with the select 30 methylators. Individual ribosomal protein hits were aligned with MAFFT (33) and concatenated. The tree was constructed using FastTree and further visualized and edited using iTOL (38, 39) to highlight methylating phyla and novel methylating groups.

We created an HMM profile of *hgcB* from experimentally verified methylators as described above for *hgcA* (12, 40) From the set of 904 putative methylators identified by *hgcA* presence, we searched for *hgcB* by using a threshold cutoff score of 70 and requiring the resulting hit to either be directly downstream of *hgcA* or within five open reading frames, as some methylators have been observed to have genes between *hgcAB*. We recognize that this criterion might be overly conservative, potentially excluding some genomes with both genes (e.g. *hgcB* on a separate contig) but prefer to avoid any false positives. This resulted in 844 of 904 identified methylators also containing *hgcB*. To create a representative HgcAB tree to compare against a concatenated ribosomal protein phylogeny, a given genome had to contain 12 or more ribosomal protein markers to ensure accurate comparisons. Additionally, we removed sequences with low bootstrap support using RogueNaRok (41). Overall, this resulted in 650 HgcAB sequences to create a representative tree.

The HgcAB protein sequences were aligned separately using MAFFT and concatenated (33). The alignment was uploaded to the Galaxy server for filtering with BMGE1.1 using the BLOSUM30 matrix, a threshold and gap rate cutoff score of 0.5, and a minimum block size of 5. (42, 43). A phylogenetic tree of the hits was constructed using RaxML with 100 rapid bootstraps (44). The tree was automatically rooted and edited using the interactive tree of life (iTOL) online tool (39). For visualization ease, we collapsed clades by the dominant, monophyletic phyla when possible. For example, if a monophyletic clade contained a majority (approximately 80-90%) of *Deltaproteobacteria* sequences and one *Nitrospirae* sequence, we collapsed and identified the clade as a whole as *Deltaproteobacteria*. This was done to broadly highlight the sparse phylogenetic nature of identified HgcAB sequences. To reduce complexity, identified HgcAB sequences of candidate phyla containing few individuals are not assigned individual colors. The full phylogenetic tree with non-collapsed branches and most resolved taxonomical names is provided in Supplementary Figure 2.

The corresponding ribosomal tree of the representative HgcAB sequences was constructed using a collection of 16 ribosomal proteins provided in the metabolisHMM package (36, 45). Each marker was searched for in each genome using specific HMM profiles for each ribosomal protein, and individually aligned using MAFFT (33). Hits for archaeal and bacterial taxa were concatenated separately to create an archaeal and bacterial ribosomal protein tree. A phylogenetic tree was constructed with RAxML with 100 bootstraps and visualized using iTOL (44, 39).

### Functional Annotations, Metabolic Reconstructions, and *in silico* PCR analysis

All genomes were functionally annotated using Prokka (46). Using the locus tag of the predicted *hgcA* protein with the above HMM-based approach, the predicted gene sequence for each *hgcA* open reading frame was obtained. *In silico* PCR of the universal *hgcA* primers and group-specific qPCR *hgcA* primers were tested using Geneious (Biomatters Ltd, Auckland, NZ), with primer sequences from Christensen et al. (20). The settings allows for both 0 and 2 mismatches, and the primer amplification “hit” results were compared to the full-genome classification of the given *hgcA* gene sequence. For genome synteny analysis of select permafrost methylators, Easyfig was used to generate pairwise BLAST results and view alignments (47). We did not use one of the *Myxococcales* MAGs as it was exactly identical to the other. The *Opitutae* genome GCA_003154355 from the UBA assembled set was not used because *hgcA* wsa found on only a very short contig.

Broad metabolic capabilities were characterized across all methylating organisms based on curated sets of metabolic HMM profiles provided in the metabolisHMM package (45, 37). An HMM of the putative transcriptional regulator was built using the alignment of the five putative transcriptional regulators with similar *hgcAB* sequences with muscle and hmmbuild (40, 48), and searched for among all methylators with hmmsearch (40). A genome was considered to have a hit for the putative transcriptional regulator if the identified protein contained a search hit e-value of greater than 1e-10 and was immediately upstream of *hgcAB* or 1-2 genes upstream. *Mer* genes for mercury transport were accessed from KofamKOALA using the corresponding threshold cutoff scores (49). Raw presence/absence results are provided in Supplementary Table 5.

Metabolic reconstruction of the MENDH-Thermoleophilia bin was supplemented with annotations using the KofamKOALA distribution of the KEGG database, and parsed for significant hits (49). The phylogeny of the MENDH-Thermoleophilia MAG was confirmed by downloading representative/reference sequences from Genbank and Refseq for the entire Actinobacteria phyla, depending on which database/designation was available for the particular class. All Thermoleophilia class genomes were downloaded as designated by the Genome Taxonomy Database (35). We included *Bacillus subtilis* strain 168 as an outgroup. The phylogeny was constructed using the alignment of concatenated ribosomal proteins as described above. The tree was constructed using RAxML with 100 rapid bootstraps and visualized with iTOL (44, 39).

### Transcriptional Activity of Methylators in a Permafrost System

To investigate the transcriptional activity of methylators in the environment, we tracked activity of the identified methylating organisms in a permafrost thaw gradient published by Woodcroft et al. 2018 (50). All 26 raw metatranscriptomes from the palsa, bog, and fen sites were downloaded from NCBI. Adapters were removed and sequences quality filtered with fastp (51). Each transcriptome was mapped to the indexed open reading frames of the 111 putative methylators identified in this ecosystem using kallisto (52). Annotated open reading frames and size in base pairs for each genome were supplied using Prokka (46). For every sample, counts were normalized by calculating transcripts per million (TPM), which equates for every 1 million reads in an RNA-seq sample, “x” amount came from a certain gene/transcript. Briefly, TPM was calculated by dividing the number of raw reads mapping to a given gene by dividing by the length of that gene in base pairs. The sum of all normalized counts was divided by 1 million to create a “per million” scaling factor. Each transcript’s normalized count is divided by the 1 million factor, which creates the final TPM value.

The *hgcA* open reading frame for each genome was identified by the corresponding locus tag from the HMM profile annotation as described above. We did not detect expression of any of the putative methylators in any of the palsa samples, and therefore did not include these samples in the results. *hgcA* expression within a phylum was calculated by total TPM expression of *hgcA* within that phylum. Total *hgcA* expression within a sample was calculated as the total TPM-normalized expression of *hgcA* counts within that sample. The total average expression of methylators within a phylum was calculated by adding all TPM-normalized counts for all genomes within a phylum, and averaged by the number of genomes within that phyla to represent the average activity of that group compared to *hgcA* activity. Expression of *hgcA* was compared to the housekeeping gene *rpoB*, which encodes the RNA polymerase beta subunit. *rpoB* loci were predicted from Prokka annotations, and we compared expression between *hgcA* and *rpoB* at the phylum level. Normalized counts of *hgcA* and *rpoB* were summed within each phylum for each sample.

### Data and Code Availability

Lake Tanganyika genomes have been deposited in Genbank under the project ID PRJNA523022 and are available at https://osf.io/pmhae/. The four Trout Bog genomes can be accessed from JGI/IMG under accession IDs 2582580680, 2582580684, 2582580694, and 2593339183. A complete workflow including all code and analysis workflows can be found at https://github.com/elizabethmcd/MEHG. All supplementary files including metadata, *hgcA* sequences and alignments, and tables are available on FigShare at https://figshare.com/account/home#/projects/70361 under the CC-BY 4.0 license.

## RESULTS AND DISCUSSION

### Identification of Diverse, Novel Putative Mercury Methylating Microorganisms

We identified nearly 1000 putative bacterial and archaeal mercury methylators spanning 30 phyla among publicly available isolate genomes and MAGs recovered from numerous environments (Figure 1). Well-known methylators among the *Deltaproteobacteria, Firmicutes, Euryarchaeota, and Bacteroidetes* represent well over half of all identified putative methylators, with the majority among the *Deltaproteobacteria* (Figure 1C). We also expanded upon groups of methylators that until recently have not been considered to be major contributors to mercury methylation. Putative methylators belonging to the *Spirochaetes* and the PVC superphylum (consisting of *Planctomycetes, Verrucomicrobia, Chlamydiae,* and *Lentisphaerae* phyla and several candidate divisions) have been identified from metagenomic contigs (13) and reconstructed MAGs from Jones et al (22). Additionally, *hgcA* sequences have been detected on metagenomic contigs belonging to the *Nitrospirae, Chloroflexi,* and *Elusimicrobia* (19, 53), but overall it is unknown how these groups contribute to methylmercury production. We identified putative methylators belonging to phyla that, to our knowledge, have not been previously described as mercury methylators. We identified 7 *Acidobacteria* methylators, 5 of which from the permafrost gradient system, in which these are considered to be the main plant biomass degraders (50). We also identified 17 *Actinobacterial* methylators, which all belong to poorly characterized lineages within the *Coriobacteria* and *Thermoleophilia* classes, as described below. A handful of putative methylators belong to recently described candidate phyla, such as Candidatus Aminicenantes, Candidatus Firestonebacteria, and WOR groups (Supplementary Figure 1D).

**Figure 1.**
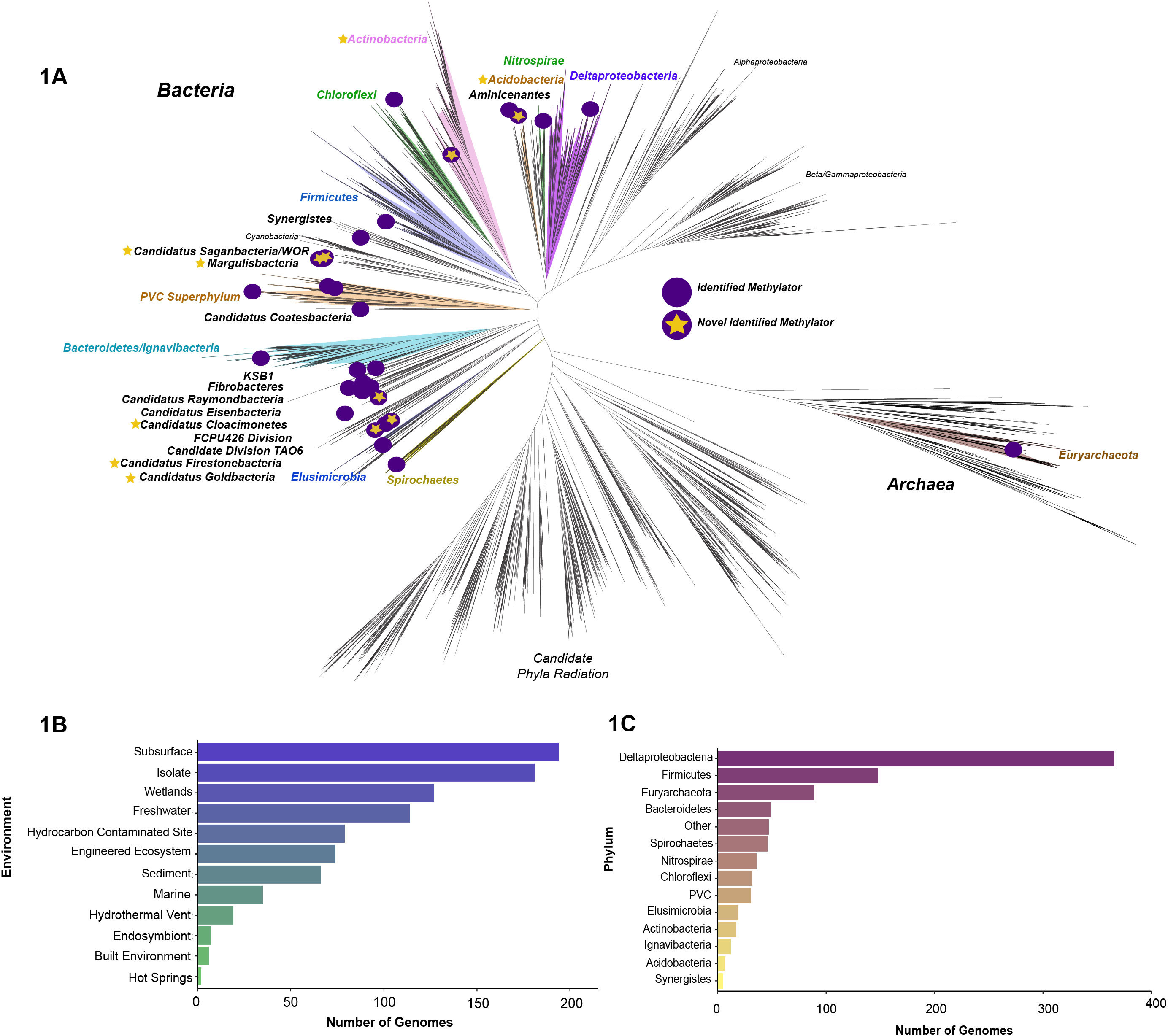
Phylogenetic Distribution and Diversity of Putative Methylators. Phylogenetic diversity and distribution of putative and novel methylators. **A)** Putative and novel methylators among the prokaryotic tree of life. The highest quality methylator from each identified phylum was selected as a representative among the tree of life. Purple dots represent methylating phyla, and purple dots with stars represent phyla that, to our knowledge, have not been identified as mercury methylators before this study. Groups in bold and/or colored are methylating groups, whereas a few uncolored groups in non-bold face are noted only for orientation. **B)** Number of putative methylating genomes found in various environments, or genomes sequenced from isolates. **C)** Number of genomes belonging to each phylum. Phylum names are given for all groups except the *Deltaproteobacterial* class of *Proteobacteria*, as no putative *Proteobacterial* methylators were identified outside of the *Deltaproteobacteria*. The group “other” contains several candidate phyla and genomes within groups with few representatives, described in Supplementary Figure 1.

We reconstructed MAGs from three freshwater lakes with diverse biogeochemical characteristics. Lake Mendota is a large, dimictic, eutrophic lake in an urban setting in Madison, WI, USA, with a sulfidic anoxic hypolimnion. Trout Bog Lake is a small, dimictic humic lake in a rural area near Minocqua, WI, that is surrounded by sphagnum moss, which leaches large amounts of organic carbon. Lake Tanganyika is a meromictic lake with an anoxic monimolimnion in the East African Rift Valley, and is the second largest lake in the world both by volume and depth, from which we recently reconstructed nearly 4000 MAGs from (23). From these reconstructed freshwater MAGs, we identified 55 putative mercury methylators. Several understudied *Verrucomicrobia* clades were recovered from the Lake Mendota hypolimnion. Members of this phylum are ubiquitous in freshwater and marine environments, display a cosmopolitan distribution, and can make up approximately 7% of the total microbial community in these ecosystems (28, 54–56). In Lake Mendota, members of the PVC superphylum account for approximately 40% of the microbial community in the hypolimnion, the majority of which do not contain *hgcA* (Peterson unpublished 2020).

A majority of the identified methylators were from subsurface aquifer system and thawing permafrost gradient assembled by Anantharaman et al. and Woodcroft et al., respectively (50, 57) (Figure 1B). Additionally, we identified methylators from hydrocarbon contaminated sites such as oil tills and sand ponds, sediments and wetlands, engineered systems, hydrothermal vents, and the built environment. A handful of methylating organisms were recovered from marine systems, as the Tara Ocean projects mostly includes samples from surface waters (58), and therefore would not include traditionally anaerobic methylating microorganisms. Interestingly, we identified marine endosymbionts of a mouthless, gutless worm (59), and Foraminifera spp. Although not as numerous as MAGs from various metagenomic surveys, we identified *hgcA* in over 150 isolate genomes, providing an expanded resource of isolates for experimentation (Supplementary Figure 1A). Using an extensive set of genomes mostly recovered from anoxic environments, we were able to greatly expand upon the known phylogenetic diversity of putative mercury methylating microorganisms.

### Novel *Actinobacterial* Methylating Lineages

To our knowledge, members of the *Actinobacteria* have never been characterized as putative methylators or found to contain the *hgcA* marker. We identified 17 *Actinobacterial* putative methylators among diverse environments, including our Trout Bog and Lake Mendota freshwater datasets, as well as within the permafrost gradient and MAGs recovered from hydrocarbon contaminated sites (50, 60). In freshwater systems, *Actinobacteria* are ubiquitous and present a cosmopolitan distribution, with specific lineages comprising up to 50% of the total microbial community (61, 62). Freshwater *Actinobacteria* traditionally fall within the class *Actinobacteria*, but have a lower abundance distribution in the hypolimnion due to decreasing oxygen concentrations (63, 64). None of the *Actinobacterial* putative methylators belong to ubiquitous freshwater lineages.

All of the 17 identified *Actinobacterial* putative methylators belong to poorly described classes or ill-defined lineages that may branch outside of the established six classes of *Actinobacteria* (62). Nine fall within the *Coriobacteriia* class, four fall within the *Thermoleophilia* class, and the other four diverge from the existing *Actinobacteria* lineages. Three of the ill-defined *Actinobacterial* MAGs have been designated within the proposed class UBA1414, with two assembled from the permafrost system and one from the Lake Mendota hypolimnion. The other ill-defined *Actinobacterial* MAG was designated within the proposed class RBG-13-55-18 and was also recovered from the permafrost system. While some *Coriobacteriia* have been identified as clinically-significant members of the human gut, all identified putative methylators belong to the OPB41 order (65), which refers to the 16S rRNA sequence identified by Hugenholtz et al. from the Obsidian Pool hot spring in Yellowstone National Park (66). Members of the OPB41 order have also been found to subsist along the subseafloor of the Baltic Sea, an extreme and nutrient poor environment (67).

Members of the *Thermoleophilia* are known as heat- and oil-loving microbes due to their growth restriction to only substrate n-alkanes (68). Within this deep-branching lineage, the two orders *Solirubrobacterales* and *Thermoleophilales* are currently recognized based on sequenced isolates and the most updated version of Bergey’s Taxonomy (69, 70). However, our *Thermoleophilia* MAGs from the hypolimnia of Lake Mendota and Trout Bog as well as two other putative *Thermoleophilia* methylators from the permafrost thawing gradient do not cluster within the recognized orders (Figure 2). According to the GTDB, these MAGs cluster within a novel order preliminarily named UBA2241, as the only MAGs recovered from this novel order to-date have been from the author’s corresponding permafrost system (35, 50). Additionally, based on collected isolates, members of the *Thermoleophilia* are assumed to be obligately aerobic (70).

**Figure 2.**
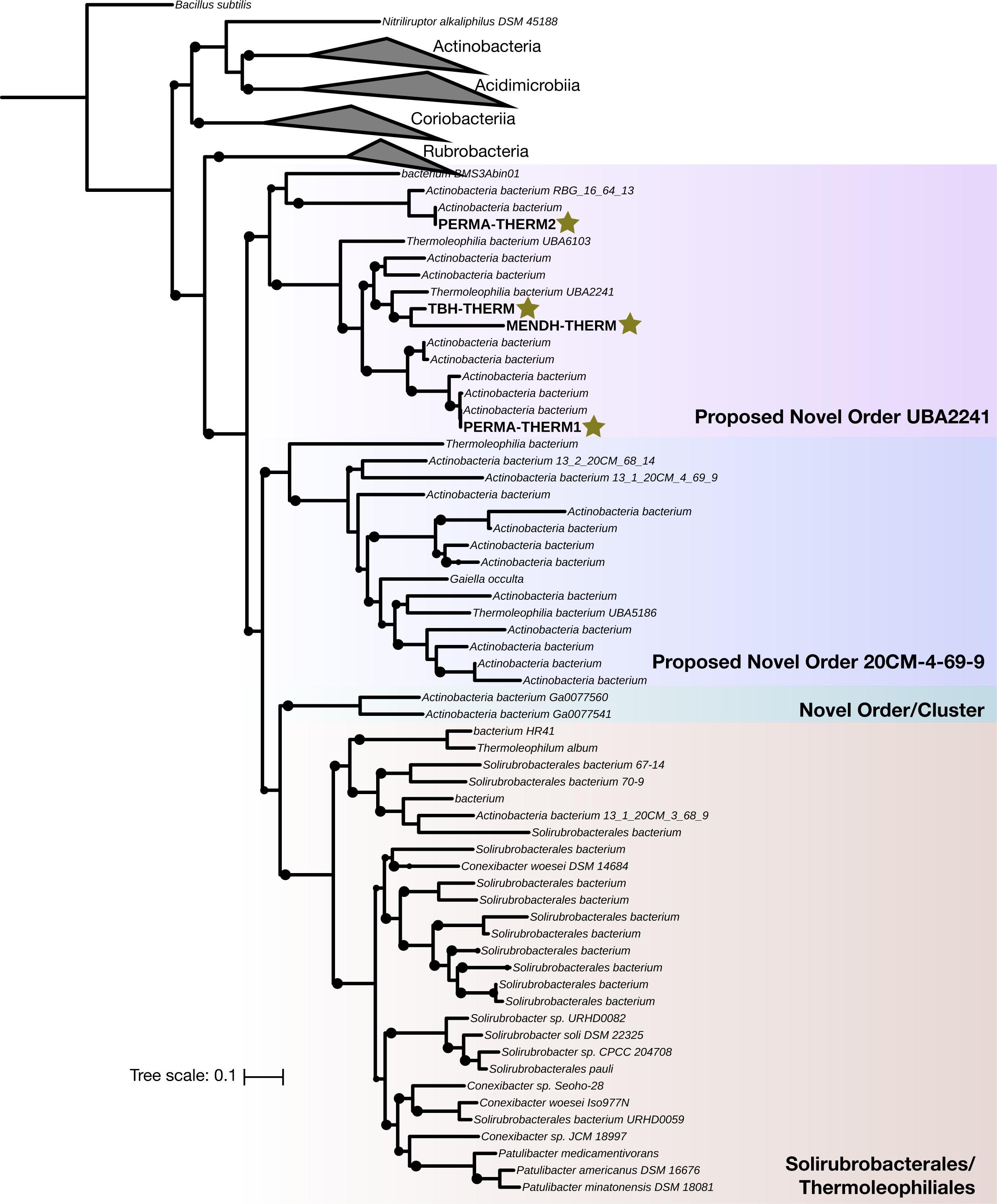
Novel Putative Methylators within the *Thermoleophilia* class of Actinobacteria. Phylogenetic tree of publicly available and assembled *Thermoleophilia* MAGs and *Actinobacteria* references. Representative or reference genomes belonging to the other 5 classes of *Actinobacteria* were used, with *Bacillus subtilis* used as an outgroup. Orders are clustered and colored by GTDB proposed designation. The phylogeny was constructed with RaxML using 100 rapid bootstraps, and nodes with bootstrap support greater than 60 are shown as black circles. Stars denote putative methylators identified in this study.

The functional potential of *Thermoleophilia* members has been poorly described due to few sequenced isolates and high-quality MAGs (71). We reconstructed the main metabolic pathways of a high-quality *Thermoleophilia* MAG, referred to as MENDH-Thermo (Supplementary Table 2). All components of the glycolytic pathway are present for using a variety of carbohydrates for growth. A partial TCA cycle for providing reducing power is present; only missing the step for converting citrate to isocitrate, which can be transported from outside sources. Interestingly, the MENDH-Thermo genome encodes the tetrahydromethanopterin-dependent *mch* and *fwdAB* enzymes, pointing to either formate utilization or detoxification (72, 73). We could not detect the presence of a putative acetate kinase, and therefore autotrophic growth through forming acetate is unlikely. Interestingly, the MENDH-Thermo genome encodes a full pathway for 4-Hydroxybutryate formation, a storage polymer (74). As the MENDH-Thermo genome can also synthesize the amino acids glycine and serine, combined with the ability to use reduced carbon compounds and form storage polymers, the MENDH-Thermo genome could exhibit a methylotrophic lifestyle, but is missing key steps for tetrahydromethanopterin cofactor synthesis and formate oxidation (75). The identification of putative methylators within the *Thermoleophilia* not only highlights novel metabolic features of putative methylators, but also of a poorly described class within the *Actinobacteria*.

### Implications for Horizontal Gene Transfer of *hgcAB*

The HgcAB protein phylogeny exhibits patterns of extensive horizontal gene transfer (HGT) compared to a concatenated phylogeny of ribosomal proteins of corresponding genomes (Figure 3). For example, *Deltaproteobacterial* HgcAB sequences are some of the most disparate sequences surveyed, clustering with HgcAB sequences of *Actinobacteria, Nitrospirae, Spirochaetes,* and members of the PVC superphylum. Although *Firmicutes* HgcAB sequences are not as disparate, they also cluster with HgcAB sequences of other phyla such as *Deltaproteobacteria* and *Spirochaetes.* The only HgcAB sequences that cluster monophyletically among our dataset are the *Bacteroidetes/Ignavibacteria* groups and *Nitrospirae*. These observations have implications for the underlying evolutionary mechanisms of the methylation pathway and modern methods for identifying methylating populations in the environment.

Gene gain/loss mediated through several HGT events have been previously suggested to underpin the sparse and divergent phylogenetic distribution of *hgcAB* (11, 22). Metabolic functions can be horizontally transferred across diverse phyla through numerous mechanisms, such as phage transduction, transposon-mediated insertion of genomic islands or plasmid exchanges, direct conjugation, and/or gene gain/loss events. We were unable to identify any *hgcA*-like sequences on any viral contigs in the NCBI database or within close proximity to any known insertion sequences or transposases, and all known *hgcA* sequences have been found on the chromosome. Structural and mechanistic similarities between the cobalamin binding domains of HgcA and the carbon monoxide dehydrogenase/acetyl-CoA synthase (CODH/ACS) of the reductive acetyl-CoA pathway (also known as the Wood-Ljungdahl pathway) suggests associations between the two pathways (11, 13). Interestingly, *hgcA* sequences of *Euryarchaeota* and *Chloroflexi* cluster together, which was recently found to also be true for the CODH of these groups (76). Inter-phylum HGT among anaerobic microorganisms has been proposed to account for approximately 35% of metabolic genes and contributes to functional redundancy across diverse phyla (77). It is plausible that a gene duplication event followed by rampant gene gain/loss and independent transfers of both the CODH and HgcA proteins could explain the disparate phylogeny, as has been respectively described for both pathways (11, 13, 76).

**Figure 3.**
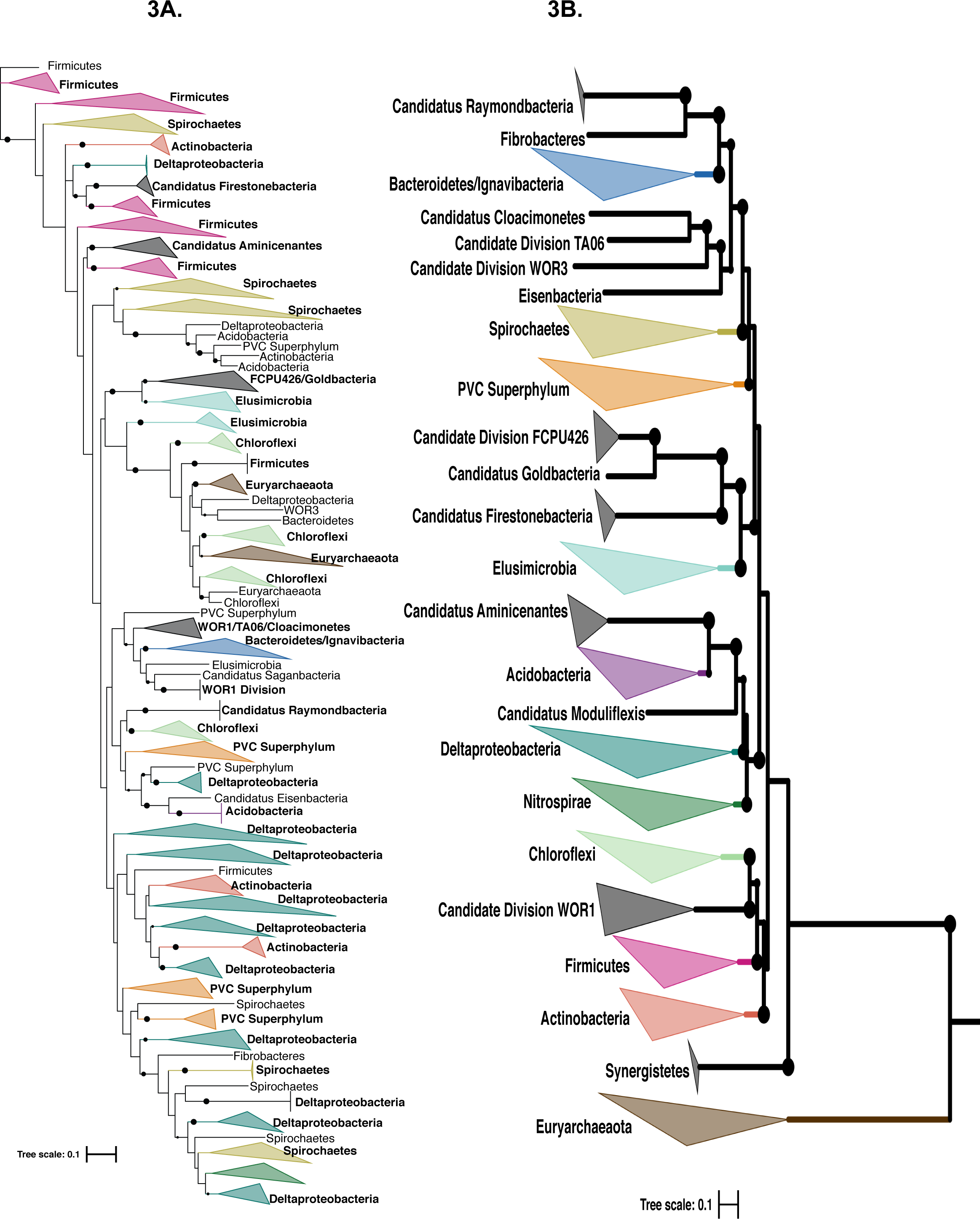
Horizontal Gene Transfer of HgcAB Relative to Corresponding Ribosomal Protein Phylogeny. Phylogeny of HgcAB compared to a concatenated ribosomal protein phylogeny of select methylators. **A)** Phylogeny of HgcAB of select methylators was constructed as described in the Materials and Methods. The tree was constructed using RaxML with 100 rapid bootstraps, and nodes with bootstrap support greater than 60 are depicted as a black circle. **B)** Corresponding concatenated ribosomal protein phylogeny of the select methylators. The tree was constructed using a set of 16 ribosomal proteins, concatenated, and built using RaxML with 100 rapid bootstraps. Nodes with bootstrap support of greater than 60 are denoted as a black circle. A fully annotated HgcAB tree is provided in Supplementary Figure 2.

To illustrate inter-phylum *hgcAB* HGT, we selected methylators assembled from a permafrost system containing *hgcAB* regions with approximately 70% DNA sequence similarity (Figure 4). Five methylators encompassing four different phyla (one *Deltaproteobacteria,* two *Acidobacteria,* one *Verrucomicrobia,* and an *Actinobacteria)* contain highly similar *hgcAB* regions, and the *hgcA* phylogeny does not match the ribosomal protein phylogeny. All *hgcAB* sequences of these methylators are preceded by a hypothetical protein that also shares sequence similarity among all methylators, but which is annotated as a putative transcriptional regulator in the *Opitutae* genome. We screened for this putative transcriptional regulator among all methylators in our dataset and identified it immediately upstream of *hgcAB* in 112 genomes (Supplementary Table 3). Interestingly, we identified 12 genomes in which the putative transcriptional regulator was upstream of *hgcAB*, but separated by one or two other genes. These genes were mostly annotated as hypothetical proteins, but included esterases, oxidoreductases, and methyltransferases. Otherwise, the gene neighborhoods of these similar permafrost *hgcAB* sequences does not share any other obvious sequence or functional similarities. The *hgcAB* genes are flanked by numerous hypothetical proteins, varying amino acid synthesis and degradation genes, and oxidoreductases for core metabolic pathways.

**Figure 4.**
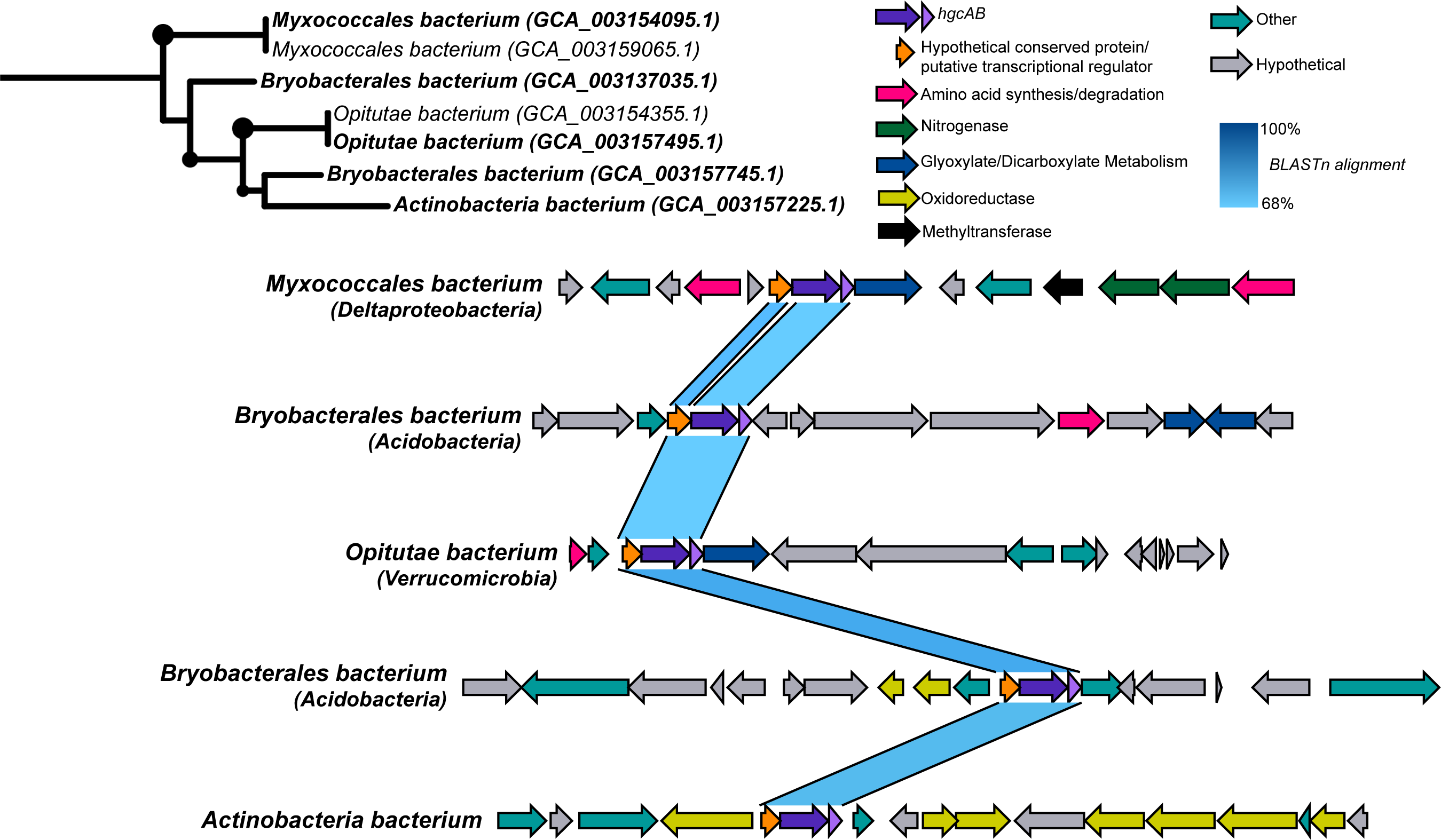
Gene Neighborhoods of Diverse Permafrost Methylators Containing similar *hgcAB* Regions. Contigs containing similar *hgcAB* regions from putative methylators identified in a permafrost thawing system (50) were compared. Pairwise nucleotide BLAST between the genes for each contig was performed and visualized in easyfig (47). Colors correspond to predicted functions of each gene. *hgcAB* genes were identified based on curated HMMs and reference sequences, while all other gene assignments were made based on Prokka predictions.

Methylating populations have been identified in the environment through a combination of nonspecific methods such as associations through 16S rRNA amplicon sequencing (15, 78, 79), *hgcA* identification on metagenomic contigs (18, 53), universal qPCR primers, and specific assays such as clade-specific qPCR primers (19, 20). However, universal and clade-specific primers were constructed based on a handful of experimentally verified methylators (20), which likely does not capture the complete or true diversity of methylating microorganisms. We tested the accuracy of these primers *in silico* on the coding regions of *hgcA* in all our identified putative methylators, using the taxonomical assignment of each putative methylator from the full genome classification (Supplementary Table 2). Depending on the number of *in silico* mismatches allowed, only experimentally verified methylators could be identified, or *hgcA* sequences belonging to other phyla would be hit (Supplementary Table 4). For example, *Deltaprobacterial-* specific *hgcA* qPCR primers picked up related *Actinobacterial, Spirochaetes,* and *Nitrospirae* sequences. Whereas the *Firmicutes*-specific primers did detect all experimentally-verified *Firmicutes* methylators and a few environmental *hgcA* sequences, overall they missed the diversity of *Firmicutes hgcA* sequences. Evidence of extensive HGT of *hgcAB* across diverse phyla suggests that “universal” or even targeted guild amplicon approaches may not be accurate. Instead, finer-scale primers of specific methylating groups could be constructed based on recovered population genomes from a specific environment containing *hgcAB* to ensure accuracy.

### Diversity of Metabolic Capabilities of Identified Methylators

We next characterized the broad metabolic capabilities among high-quality isolates and MAGs. From our analysis of metabolic profiles spanning several biogeochemical cycles (37), *hgcA* is the only genetic marker linking high-quality methylating genomes other than universally conserved markers (Figure 5). We detected putative methylators with methanogenesis and sulfate reduction pathways, as is expected for archaeal and bacterial methylating guilds, respectively (17, 22). Although the presence of sulfate has been thought to select for methylating bacteria and drive methylation, very few guilds we examined exhibited dissimilatory sulfate reduction pathways. We detected machinery for dissimilatory sulfate reduction within the *Acidobacteria, Deltaproteobacteria, Firmicutes,* and *Nitrospirae*. However, for example, only half of the *Deltaproteobacterial* methylators screened contained the *dsrABD* subunits for sulfate reduction. Similarly, we detected few guilds with the sulfate adenylyltransferase (*sat)* marker for assimilatory sulfate reduction for incorporating sulfur into amino acids. This suggests that mercury methylating groups may be composed of guilds other than canonical SRBs and methanogenic archaea.

**Figure 5.**
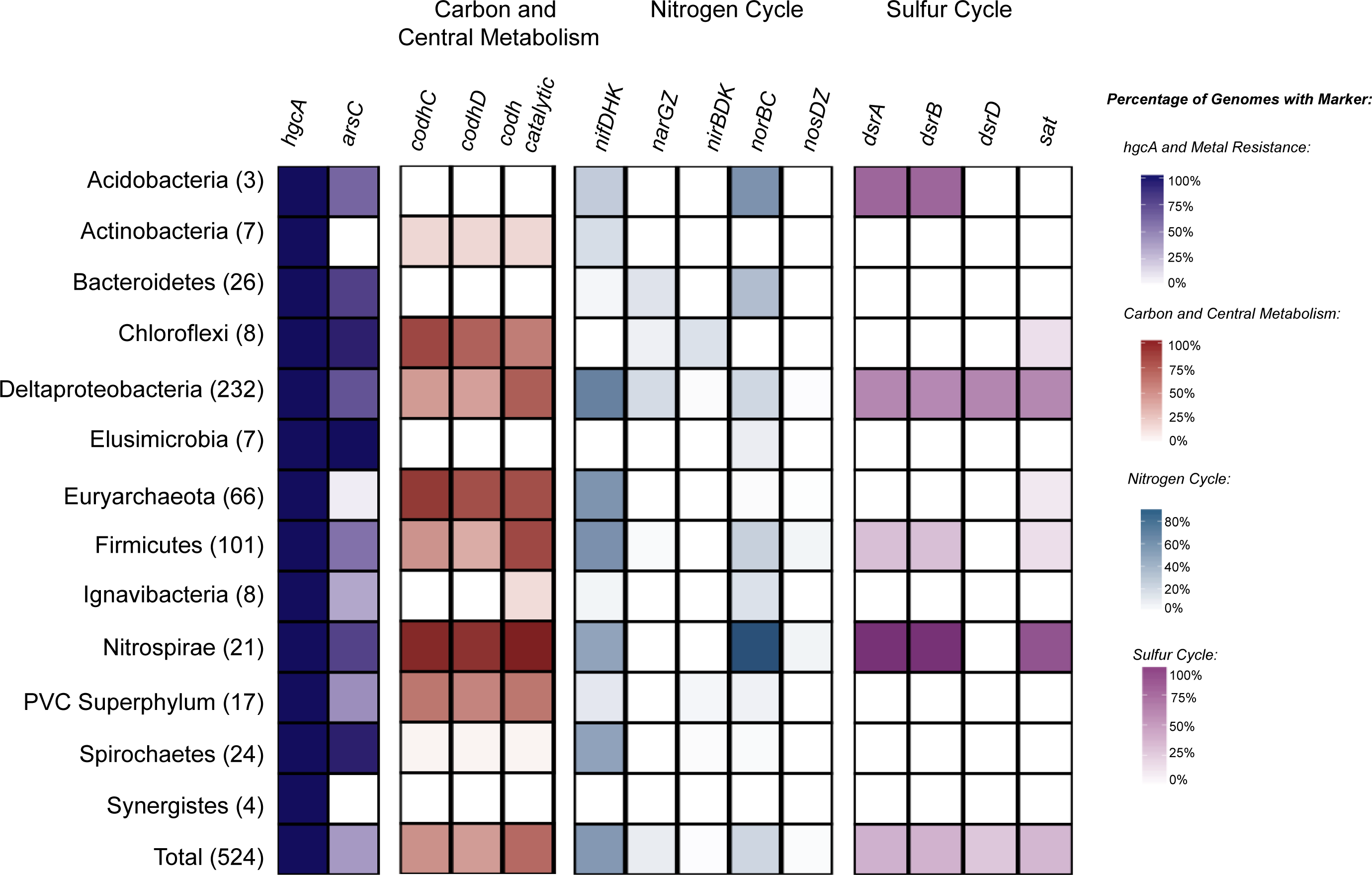
Metabolic Characteristics of Putative Methylators. Summary of broad metabolic characteristics present among high-quality putative methylators. Genomes with > 90% completeness and <10% redundancy and belonging to phyla with more than 5 genomes in that group were screened. HMM profiles spanning the sulfur, nitrogen, and carbon cycles, and metal resistance markers were searched for. The intensity of shading within a cell equates to the corresponding colored legend of percentage of genomes within that group that contain that marker. *hgcA* = Mercury methylation corrinoid protein, *arsC* = arsenite reductase, *codhCD/catalytic* = carbon monoxide dehydrogenase C,D, and catalytic subunits, *nifDHK* = nitrogenase subunits, *narGZ* = nitrate reductase subunits, *nirBDK* = nitrite reductase subunits, *norBD* = nitric oxide reductase subunits, *nosDZ* = nitrous oxide reductase subunits, *dsrABD* = dissimilatory sulfite reductase, *sat* = sulfate adenylyltransferase. Individual *nif, nar, nir, nor,* and *nos* subunit presence/absence results were averaged together, respectively.

Interestingly, we found that several methylating guilds contain the essential subunits that form the nitrogenase complex for nitrogen fixation. The three main components of the nitrogenase complex are *nifH* and *nifD/nifK*, which encode the essential iron and molybdenum-iron protein subunits, respectively. We detected machinery for the nitrogenase complex within the *Acidobacteria, Actinobacteria, Deltaproteobacteria, Euryarchaeota, Firmicutes, Nitrospirae,* and *Spirochaetes* methylators. Notably, over half of all *Deltaproteobacterial* methylators and approximately half of all *Firmicutes* methylators contain all three subunits for the nitrogenase complex. Recently, the presence of nitrate has been suggested to regulate methylmercury concentrations (80) leading to nitrate addition in an attempt to control methylmercury accumulation in freshwater lakes (81). Although we detected parts of the denitrification pathway in a handful of lineages, none of the putative methylators contained a full denitrification pathway for fully reducing nitrate to nitrogen gas. Previous studies have identified putative methylators among *Nitrospina* nitrite oxidizers in marine systems (18, 82). Although we did not detect putative *Nitrospina* methylators (possibly due to these *hgcA* sequences being unbinned, within low-quality MAGs, or low-confidence hits), links between the nitrogen cycle and mercury methylation warrants further exploration.

As mentioned above, structural and functional similarities between HgcA and the carbon monoxide dehydrogenase/acetyl-CoA synthase (CODH/ACS) suggests associations between the two pathways. This also raises the question if methylators use the reductive acetyl-CoA pathway for autotrophic growth. We screened for the presence of the codhC and codhD subunits of the carbon monoxide dehydrogenase, which have been suggested as potential paralogous origins of HgcA (11, 13). Among high-quality putative methylators, we detected both subunits among *Actinobacteria, Chloroflexi, Deltaproteobacteria, Euryarchaeota, Firmicutes, Nitrospirae,* and members of the PVC Superphylum. We also screened for the presence of the CODH catalytic subunit, which would infer acetyl-CoA synthesis. Although we detected the CODH catalytic subunit, it was not detected in all methylating lineages, such as the *Elusimicrobia, Synergistes, Spirochaetes.* This suggests other growth strategies than autotrophic carbon fixation among a large proportion of methylators, such as fermentation pathways as suggested previously by Jones et al. (22).

Interestingly, over 50% of the putative methylators contain a thioredoxin dependent arsenate reductase, which detoxifies arsenic through the reduction of arsenate to arsenite (83, 84). As(V) reduction to the even more toxic As(III) is tightly coupled to export from the cell, which has interesting similarities to the methylation system. Hg(II) uptake has been shown to be energy dependent and therefore is imported through active transport mechanisms (85). Hg(II) uptake is highly coupled to MeHg export, suggesting the methylation system may act as a detoxification mechanism against environmental Hg(II) (85). Interestingly, nearly none of the screened putative methylators contain the canonical *mer* mercury resistance genes, including the *merA* mercury reductase to detoxify Hg(II) to Hg(0). Additionally, very few members of the *Deltaproteobacteria, Euryarchaeota,* and *Firmicutes* contain the *merP* periplasmic mercuric transporter, suggesting a different mechanism for mercury transport and detoxification than the *mer* system. Overall, putative methylators are composed of metabolic guilds other than the classical SRBs and methanogenic archaea and employ diverse growth strategies other than autotrophic carbon fixation.

### Expression of Putative Methylators in a Permafrost System

While the connection between the presence of the *hgcA* marker and methylation status has been well established in cultured isolates, there have been few studies investigating *hgcA* transcriptional activity under laboratory conditions (86) or in a specific environment (87). Additionally, the mere presence of a gene does not predict its functional dynamics or activity in the environment. Assessing actively contributing microbial populations to methylmercury production is important for identifying constraints on this process. Genome-resolved metatranscriptomics provides an ideal approach for exploring methylating population activity in the natural environment. Woodcroft et al. sampled across three sites of the Stordalen Mire peatland in northern Sweden (50). This site includes well-drained but mostly intact palsas, intermediately-thawed bogs with *Sphagnum* moss, and completely thawed fens. Extensive spatiotemporal sampling paired with genome-resolved metagenomic, metatranscriptomic, and metaproteomic sequencing was applied to understand microbial contributions to carbon cycling along the thawing gradient. We identified 111 putative methylators among MAGs assembled from this permafrost system spanning well-known methylators belonging to the *Deltaproteobacteria, Firmicutes*, and *Euryarchaeota*, but also less explored putative methylators within the *Acidobacteria, Actinobacteria, Elusimicrobia*, and *Verrucomicrobia*. We chose this extensive dataset to explore gene expression patterns of putative methylators since it is largely an anaerobic environment, and has potential links for methylmercury production due to the buildup of mercury gas beneath frozen permafrost layers (88, 89).

We mapped the transcriptional reads for each sample to the open reading frames of all identified putative methylators, normalized by transcripts per million (TPM). We specifically investigated the depth-discrete expression patterns of *hgcA* within phylum-level groups (Figure 6). We were unable to detect *hgcA* transcripts at any time-point or depth among the palsa sites or the shallow bog depths, and therefore these sites and depths are not included. The highest total *hgcA* transcript abundance was detected in the fen sites, which is the most thawed along the permafrost gradient. The highest expression of *hgcA* was contributed from the *Bacteroidetes* in the medium and deep depths of the fen sites across multiple time-points, followed by those belonging to *Deltaproteobacteria* in the medium and deep depths of the bog sites. Interestingly, *hgcA* expression in the bog samples are almost exclusively dominated by *Deltaproteobacteria*, whereas the fen samples contain a more diverse collection of phyla expressing *hgcA*, albeit *Bacteroidetes* is the most dominant. Overall, methylators belonging to *Bacteroidetes* and *Deltaproteobacteria* also constitute the most highly transcriptionally active members across all samples.

**Figure 6.**
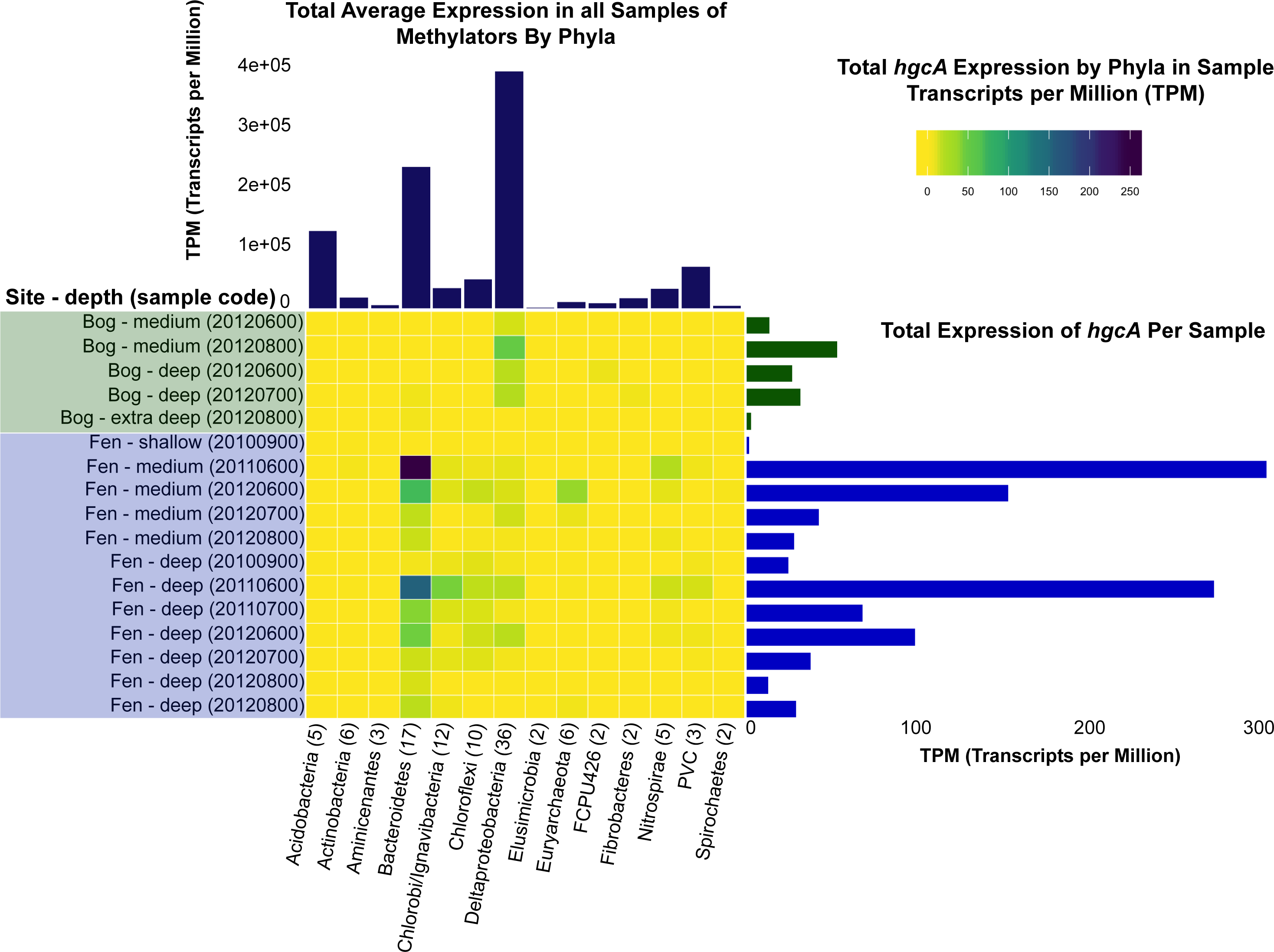
Transcriptional Activity of Putative Methylators in a Permafrost Thawing Gradient. Mapped metatranscriptomic reads to 111 putative methylators identified in a permafrost thawing gradient (50). Bog and fen samples are shown, as no expression of *hgcA* was detected in any of the palsa samples. **Center)** Total reads mapping to the *hgcA* gene within a phylum per sample normalized by transcripts per million (TPM). **Right)** Total *hgcA* counts per sample. **Top)** Average overall expression of genomes within a phylum across all samples.

All of the putative *Bacteroidetes* methylators transcribing *hgcA* belong to the vadinHA17 class. The name vadinHA17 was given to the 16S rRNA clone recovered from an anerobic digester treating winery wastewater, referring to the vinasses anerobic digestor of Narbonne (VADIN) (90). Members of the vadinHA17 group remain uncultured, and have been suggested to be involved in hydrolyzing and degrading complex organic matter, co-occurring with methanogenic archaea (91, 92). Currently, the three publicly available *Bacteroidetes* strains containing *hgcA* in belong to the *Paludibacteraceae* and *Marinilabiliaceae*, and are not closely related to members of the vadinHA17 class. However, the dominant *Deltaproteobacterial* member expressing *hgcA* in the bog samples is classified within the *Syntrophobacterales* order, specifically a member of *Smithella spp*. Among the publicly available sequenced Deltaproteobacterial isolates we identified *hgcA* in, 3 are among the *Syntrophobacterales* order, including the *Smithella* sp. F21 strain. This methylating strain may be an appealing experimental system to further explore *hgcA* expression patterns under different conditions.

We compared *hgcA* transcript abundance against that of the housekeeping gene *rpoB,* which encodes the RNA polymerase beta subunit, to differentiate *hgcA* activity against overall expression (Supplementary Figure 3). Low levels of *hgcA* expression were detected within the *Ignavibacteria, Chloroflexi, Euryarchaeota*, *Nitrospirae*, and *Planctomycetes*. Within the *Ignavibacteria* and *Chloroflexi*, expression of *rpoB* is on average higher in most samples than that of *hgcA.* This same trend occurs within the *Deltaproteobacteria*, where on average *rpoB* expression is higher than that of *hgcA*, although the majority of total *hgcA* transcripts within the bog sites is contributed to *Deltaproteobacterial* methylators. We did not detect *hgcA* transcripts within the *Aminicenantes* and *Elusimicrobia*, but genomes within these groups exhibited low activity overall as shown by their *rpoB* expression. Interestingly, we found examples of genomes within phyla that exhibited appreciable amounts of overall transcriptional activity, but did not detect *hgcA* expression. We did not detect *hgcA* expression within the *Acidobacteria* or *Actinobacteria*, but did detect *rpoB* expression. These groups exhibited higher overall expression than groups in which we detected *hgcA* expression, such as the *Nitrospirae*.

This may have interesting implications for understanding methylmercury production in the environment, as currently the presence of *hgcAB* is used to connect specific microorganisms to biogeochemical characteristics. However, it may be possible that only a few groups exhibit *hgcA* activity and actually contribute to methylmercury production. Although this study does not contain environmental data on inorganic mercury or methylmercury concentrations, these results provide insights into *hgcA* expression for further follow-up.

## CONCLUSIONS

In this study, we greatly expanded upon the known diversity of microorganisms that perform mercury methylation. We demonstrated using a set of publicly available isolate genomes, MAGs, and novel freshwater MAGs that putative methylators encompass 30 phylum-level lineages and diverse metabolic guilds. Apparent extensive HGT of the diverse *hgcAB* region poses unique challenges for screening methylating populations in the environment, in which “universal” amplicon or even group-specific approaches may not accurately reflect the true phylogenetic origin of specific methylators. Furthermore, genome-resolved metatranscriptomics of putative methylators in a thawing permafrost system revealed that specific methylating populations are transcriptionally active at different sites and depths. These results highlight the importance of moving beyond identifying the mere presence of a gene to infer a specific function and investigating transcriptionally active populations. Overall, genome-resolved ‘omics techniques are an appealing approach for accurately assessing the controls and constraints of microbial methylmercury production in the environment.

## ACKNOWLEDGEMENTS

We would like to thank the many authors and groups for making a vast and valuable amount of genomic, metagenomic, and metatranscriptomic data publicly available, as our work was greatly improved by these datasets. We thank the U.S. National Science Foundation North Temperate Lakes Long-Term Ecological Research site (NTL-LTER DEB-1440297) for providing funding of the Microbial Observatory for long-term sampling of Lake Mendota and Trout Bog. Funding was also provided to KDM by the Wisconsin Alumni Research Foundation. We thank members of the McMahon lab for constructive feedback on drafts of the manuscript, specifically Amber White. We thank Kristopher Kieft and Adam Breister for providing NCBI viral searches and tree of life ribosomal sequences, respectively. We thank the U.S. Department of Energy Joint Genome Institute for sequencing and assembly (CSPs 394 and 2796). This research was performed in part using the computer resources and assistance of the Center for High-Throughput Computing (CHTC) at UW-Madison in the Department of Computer Sciences. CHTC is supported by UW-Madison, the Advanced Computing Initiative, the Wisconsin Alumni Research Foundation, the Wisconsin Institute for Discovery, and the National Science Foundation. CHTC is an active member of the Open Science Grid, which is supported by the National Science Foundation and the U.S. Department of Energy Office of Science. Virtual machines were also accessed for computing through UW-Madison’s Campus Computing Infrastructure. This research was also performed in part using the Wisconsin Energy Institute computing cluster, which is supported by the Great Lakes Bioenergy Research Center as part of the U.S. Department of Energy Office of Science.

## FIGURE LEGENDS

**Supplementary Figure 1.**
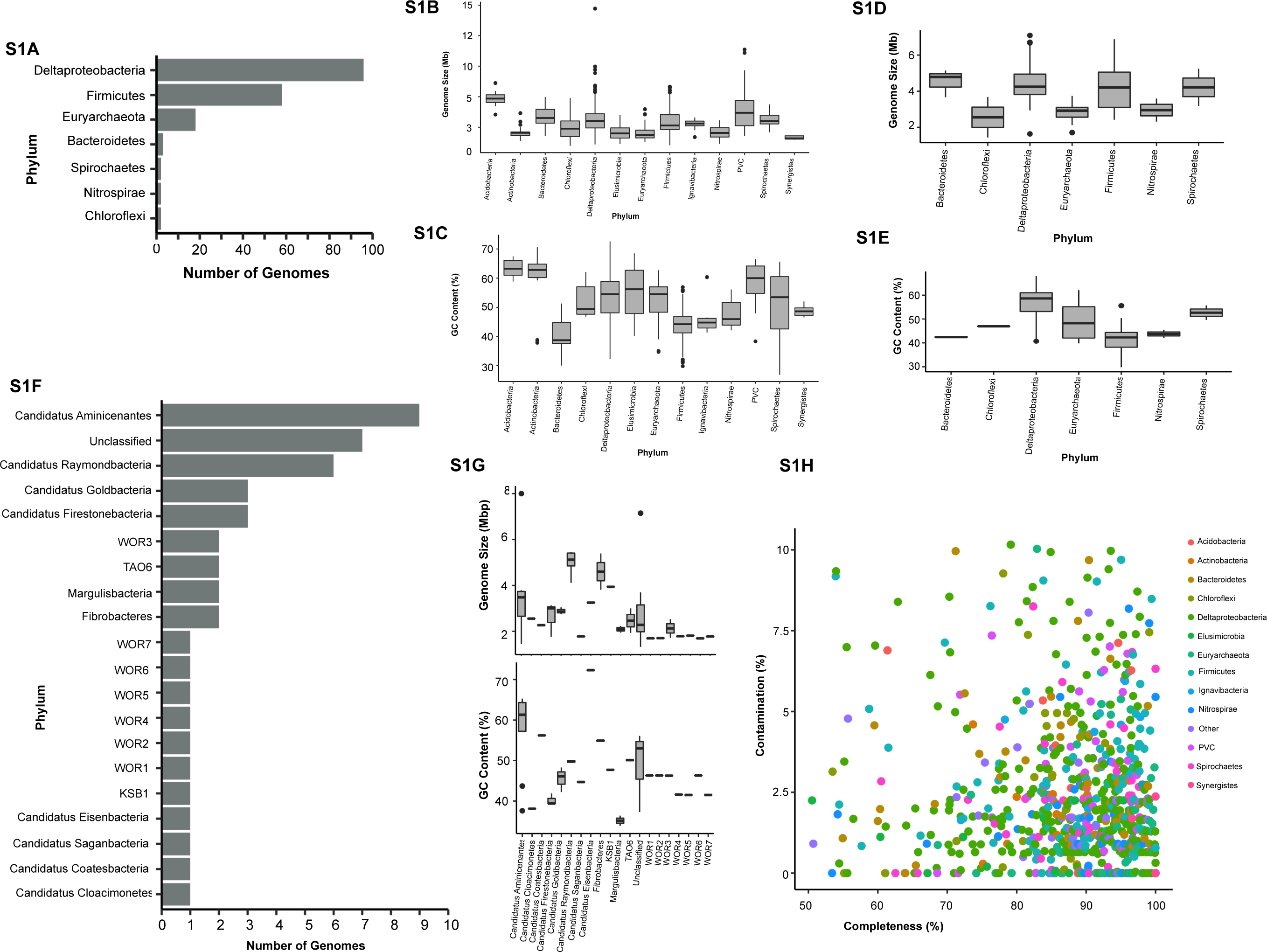
Overall Putative Methylating Genome and MAG Statistics. **A)** Number of putative methylators that are isolates and the phylum they belong to. **B)** Estimated genome size of all putative methylators. **C)** Estimated GC content of all putative methylators **D)** Estimated genome size of putative methylating isolates. **E)** Estimated GC content of putative methylating isolates. **F)** Number of genomes belonging to “other” phyla designated in Figure 1B. **G)** Estimated genome size and GC content of various candidate phyla methylators. **H)** Estimated genome completeness and contamination of putative methylators that are MAGs. Only medium quality (>50% complete and <10% redundancy) putative methylating MAGs were included in the dataset.

Supplementary Figure 2. Full Annotated Reference HgcAB tree Reference HgcAB tree of select methylators with uncollapsed nodes showing individual branch labels. Branch labels are sourced either from the name of the Genbank assembly, or custom name given to newly assembled Lake MAGs. Corresponding assembly names, classifications, and genome characteristics are provided in Supplementary Table 1. Available at https://figshare.com/account/projects/70361/articles/11620356

**Supplementary Figure 3.**
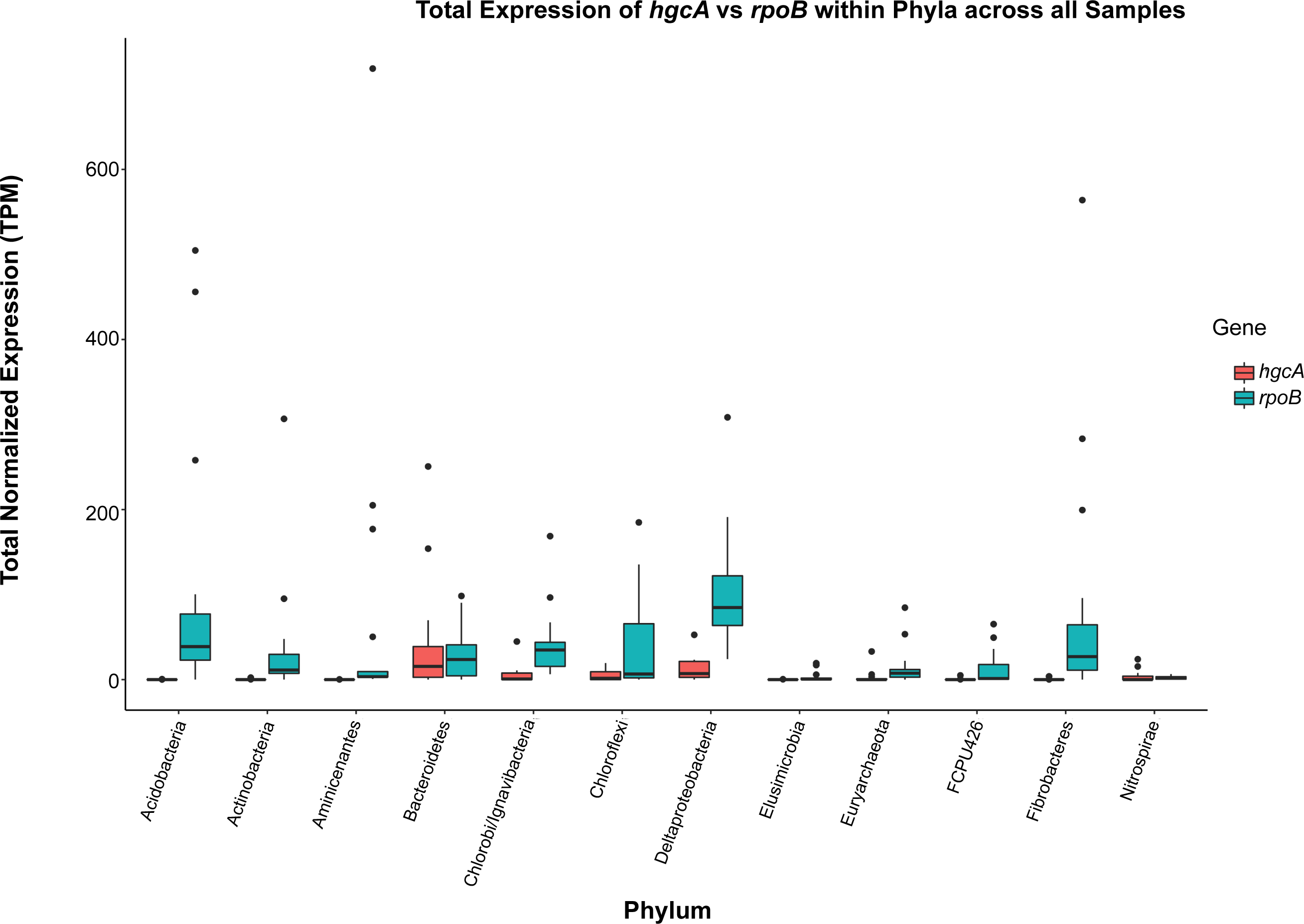
Expression of *hgcA* compared to housekeeping gene rpoB. Comparison of expression of *hgcA* and the beta subunit of the RNA polymerase *rpoB* of putative methylators in the permafrost thawing gradient. Each data point is the sum of normalized counts by transcripts per million (TPM) within a sample for that phylum. Therefore all points of the boxplot are comparing total normalized counts of gene expression between different samples within a phylum. Expression of samples within palsa sites are not included since we did not detect any *hgcA* expression in any of these samples, and did not include these results in Figure 6.

## SUPPLEMENTARY TABLES AND DATA FILES

*All supplementary files are available at* https://figshare.com/projects/Expanded_Diversity_and_Metabolic_Flexibility_of_Microbial_Mercury_Methylation/70361.

**Supplementary Table 1. Genome Characteristics** Genome characteristics for all Genbank accessions and assembled freshwater MAGs containing confident *hgcA* hits. Available at https://figshare.com/articles/mehg-metadata/10062413

**Supplementary Table 2. MENDH-Thermo Metabolic Reconstruction Annotations** Annotation results for specific pathways in the MENDH-Thermo MAG performed with Prokka and KofamKOALA. Available at https://figshare.com/account/projects/70361/articles/11620104

**Supplementary Table 3. Putative Conserved *hgcAB* Regulator Results** Results for searching for putative conserved regulator of *hgcAB* found in permafrost methylators. Available at https://figshare.com/account/projects/70361/articles/11620107

**Supplementary Table 4. Universal and Group-Specific *hgcA* Primer Results** *In silco* PCR results of universal and group-specific *hgcA* primer sets. The ORNL universal, Archaeal-specific, Deltaproteobacterial-specific, and Firmicutes-specific primer sets were ran in Genious allowing for both 0 and 2 mismatches, and matches with the taxonomical name in the metadata in Supplementary Table 1. Available at https://figshare.com/account/projects/70361/articles/10093535

**Supplementary Table 5. Presence/Absence Matrix of Metabolic Markers Among High-Quality Methylators** Raw presence/absence matrix of select metabolic markers without the conversion of results greater than 1 converted to 1, due to uncertainty of copy number variation in MAGs. Available at https://figshare.com/account/projects/70361/articles/10203530

Data File 1. HgcA Hidden Markov Profile

*Available at* https://figshare.com/account/projects/70361/articles/10084610

Data File 2. FASTA Formatted file of all Identified HgcA Protein Sequences

*Available at* https://figshare.com/account/projects/70361/articles/10084601

Data File 3. HgcB Hidden Markov Profile

*Available at* https://figshare.com/account/projects/70361/articles/11620113

Data File 4: HgcAB Alignment

*Available at* https://figshare.com/account/projects/70361/articles/11620116

## Notes

https://figshare.com/projects/Expanded_Diversity_and_Metabolic_Flexibility_of_Microbial_Mercury_Methylation/70361

